# Informative Missingness in Nominal Data: A Graph-Theoretic Approach to Revealing Hidden Structure

**DOI:** 10.1101/2025.08.22.670516

**Authors:** Ehsan Zangene, Veit Schwämmle, Mohieddin Jafari

## Abstract

Missing data is often treated as a nuisance, routinely imputed or excluded from statistical analyses, especially in nominal datasets where its structure cannot be easily modeled. However, the form of missingness itself can reveal hidden relationships, substructures, and biological or operational constraints within a dataset. In this study, we present a graph-theoretic approach that reinterprets missing values not as gaps to be filled, but as informative signals. By representing nominal variables as nodes and encoding observed or missing associations as edges, we construct both weighted and unweighted bipartite graphs to analyze modularity, nestedness, and projection-based similarities. This framework enables downstream clustering and structural characterization of nominal data based on the topology of observed and missing associations; edge prediction via multiple imputation strategies is included as an optional downstream analysis to evaluate how well inferred values preserve the structure identified in the non-missing data. Across a series of biological, ecological, and social case studies, including proteomics data, the BeatAML drug screening dataset, ecological pollination networks, and HR analytics, we demonstrate that the structure of missing values can be highly informative. These configurations often reflect meaningful constraints and latent substructures, providing signals that help distinguish between data missing at random and not at random. When analyzed with appropriate graph-based tools, these patterns can be leveraged to improve the structural understanding of data and provide complementary signals for downstream tasks such as clustering and similarity analysis. Our findings support a conceptual shift: missing values are not merely analytical obstacles but valuable sources of insight that, when properly modeled, can enrich our understanding of complex nominal systems across domains.

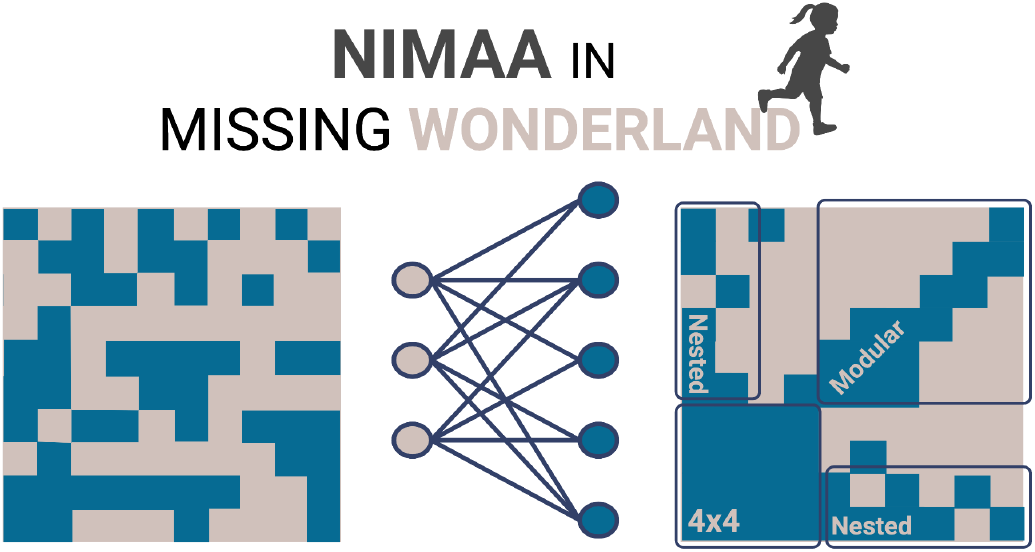

Shiny app address https://ehsan-zangene.shinyapps.io/nimaa_app/

## 1. Introduction

Missing values in datasets are often considered problematic artifacts that need to be corrected or removed through imputation ^1–3^. This approach can be effective, especially when we understand and can distinguish between missing at random (MAR) and missing not at random (MNAR) values ^1,2,4^. However, a careful examination of missingness patterns or forms can itself provide meaningful structural insights ^5,6^. This is particularly important for nominal data, which consists of multiple categorical labels without inherent numeric or ordinal relationships.

Such data, prevalent acros biomedicine, social sciences, and ecology ^7–9^ classify observation into discrete groups (e.g., patient IDs, drug names, protein accession, cell types, and species) and, particularly in cancer research, have been leveraged to uncover novel dependencies and therapeutic strategies^10^. are used to classify observations into discrete groups, such as patient IDs, drug names, protein ACs, cell types, or species to find novel dependencies and therapies in different situraitons for example in case of cancer reserach. Because nominal data lack intrinsic order or magnitude, conventional statistical methods ^11,12^ are limited in their ability to analyze them directly, and missing values pose additional challenges for interpretation and analysis. In other words, the common methods overlook the potentially informative form of missing values, which may reflect underlying biological variability, data collection constraints, or system-level organization. This limitation highlights a critical need for frameworks that can extract information from the structure and form of observed and missing nominal label associations without bearing assumptions about the causes of missingness.

The idea that missing values can carry information has been explored previously, particularly in the context of time-series data and supervised learning frameworks, where informative missingness is modeled as an explicit feature for prediction. For example, missing patterns have been leveraged to improve classification performance in longitudinal datasets^13^. In contrast, NIMAA focuses on nominal data, where labels lack intrinsic order or magnitude and standard feature-based modeling is not directly applicable. Our contribution lies in a graph-theoretic treatment of missingness, where the topology and arrangement of absent associations in bipartite networks are analyzed directly, without relying on supervised labels or temporal structure. The growing interest in network-based modeling, particularly within precision medicine, has led to the development of numerous tools for analyzing complex interactions among biological entities ^14,15^. Here, we develop a nominal data mining approach based on bipartite network modeling to explore hidden present and missing relationships among the nominal labels. Using nine datasets spanning biological and social domains, we demonstrate how patterns of missingness can influence the interpretation of the data and the underlying systems they represent. We also present this framework in the NIMAA R Shiny application built on the NIMAA R package ^16^, which provides a reproducible and interactive pipeline for applying this methodology. The major functions of the NIMAA pipeline are summarized in the overview flowchart shown in **Fig. 1**.

**Figure 1.**
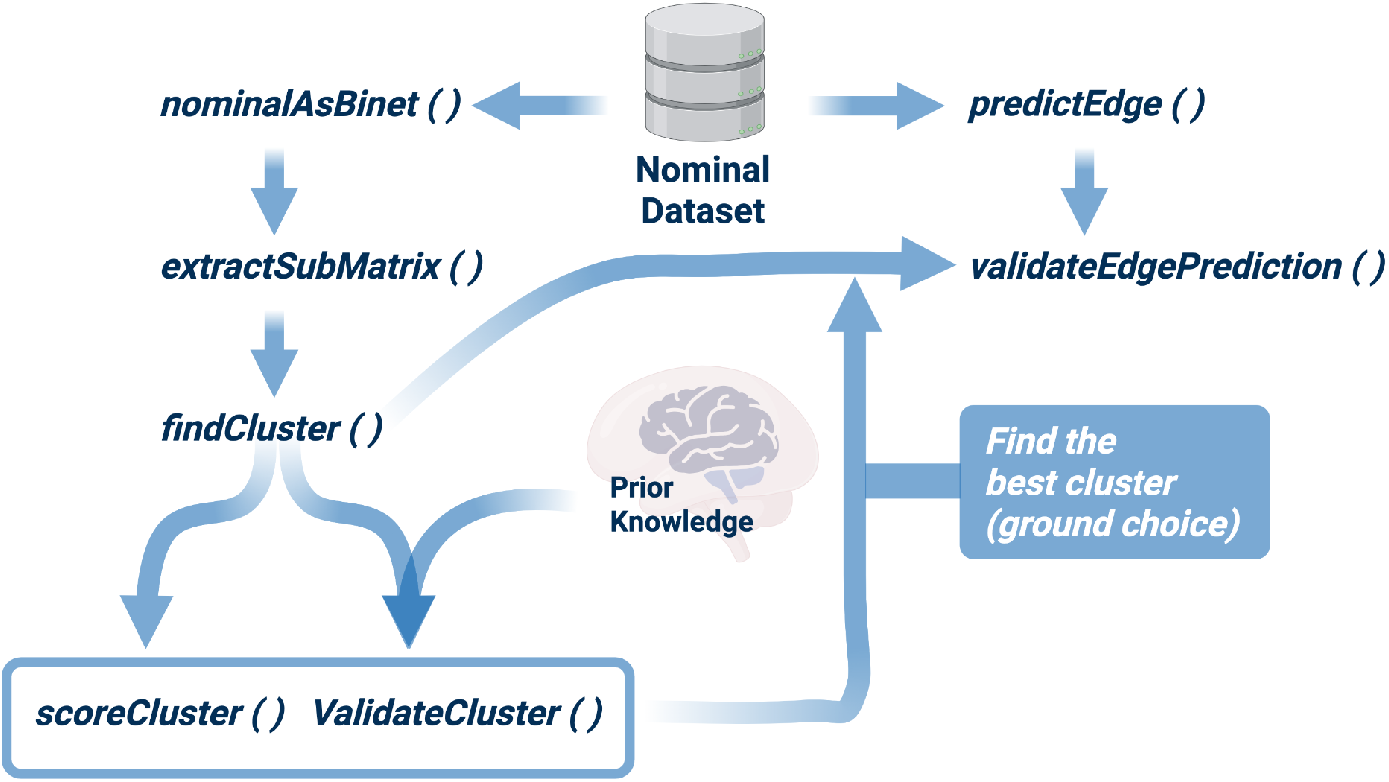
Overview flowchart of the NIMAA pipeline providing major functions. The plotIncMatrix() function displays information about the incidence matrix derived from input data, including dimensions, the proportion of missing values, and the matrix image. It also returns the incidence matrix object. In the case of a weighted bipartite network, it is recommended to use the dataset with the fewest missing values for subsequent steps. The extractSubMatrix() function extracts submatrices that contain non-missing values. The findCluster() function performs seven commonly used network clustering algorithms. Additionally, it offers the option to preprocess the input incidence matrix by projecting the bipartite network into unipartite networks. The scoreCluster() function calculates internal cluster validity measures such as entropy and coverage. The scoring is based on the weight fraction of intra-cluster edges compared to the total weight of all edges in the network. The validateCluster() function assesses the similarity of a given clustering method to a provided ground truth. It computes external cluster validity measures, including *corrected*.*rand* and *Jaccard* similarity. The predictEdge() function employs specified imputation methods to fill in missing values in the input data matrix. The output is a list where each element represents an updated version of the input data matrix without any missing values, generated using a separate imputation method. The validateEdgePrediction() function compares the clustering results after imputation with a predefined benchmark to evaluate their similarity. The benchmark is established in the previous step by validating clustering on submatrices with non-missing values. This function performs the analysis for multiple user-specified imputation methods.

## 2. Methods and Implementation

To provide a theoretical overview of our approach to nominal data mining using the NIMAA pipeline, consider a data set 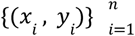 where *x*_*i*_ ∈ *V and y*_*i*_ ∈ *W* are nominal variables. This data set can be represented as a bipartite network in which *V* and *W* are disjoint node sets of the network, and each observation (*x*_*i*_, *y*_*i*_) represents a link between corresponding nodes. If each link has an associated measurement, the network becomes weighted. The connectivity structure is represented by a bi-adjacency (or incidence) matrix *A*. Two projections of the network can be taken, one for *V* and one for *W*. The adjacency matrix for the projection on part *V* is *A* × *A*^*T*^ where *A*^*T*^ is the transpose of the matrix *A*. The adjacency matrix of the projection on part *W* can also be defined using *A* × *A*^*T*^. This projection approach allows us to implement network analysis techniques for the unipartite to the bipartite case. Since these projections capture pairwise similarities within each node set, community detection on the projected networks reveals groups of similar nodes.

Beyond basic similarity and clustering analysis, bipartite networks often exhibit higher-order structural patterns. Two key examples are nestedness and modularity. Nestedness, defined as the tendency for nodes to interact with subsets of the interaction partners of better-connected nodes, is one of the important structural patterns identified in bipartite networks ^17^. Nestedness describes a hierarchical ordering of nodes in which more specialized nodes interact with a subset of the partners with whom more generalized nodes interact. It is common to rank rows and columns of the incidence matrix to visualize nestedness. As a result, the matrix, which is typically displayed in the lattice format of filled and empty squares, presents a decreasing connectivity with a large number of connections packed in the upper left corner. The most popular nestedness estimator is the “temperature” of Atmar and Patterson ^18^. A median line in the ranked incidence matrix is used to create this index. This line divides the matrix into holes and full cells equally (zeros and ones). The “temperature” index quantifies the dispersion of holes in relation to the median. In contrast to nestedness, which reflects hierarchical interaction patterns, modularity captures the presence of distinct communities within the network. Modularity identifies a network’s community structure as distinct clusters of interactions, with more connections within communities than between communities ^19^. To ensure meaningful structural analysis, we first compute nestedness and modularity indices and then address missing data by identifying and isolating complete submatrices within the incidence matrix. We might be interested in extracting the non-missing part of a weighted bipartite network. In this case, we rearrange the incidence matrix to find an approximation of the complete submatrix that includes non-missing values. It should be noted that for a given matrix, it is NP-hard to find the ‘largest’ column- or row-wise submatrix without any missing value ^20^.

Additionally, when complete submatrices are insufficient for downstream analysis, we apply imputation methods—such as mean, median, and correspondence analysis (CA)—to estimate missing values. These imputed values enable us to generate optimized network projections (see below). These networks show the similarity between each node in each part of the bipartite network. Clustering analysis is used to explore the structural pattern of similarities between nodes. When we have prior knowledge about nodes, we compare clustering results with node labels based on prior knowledge. The Jaccard and Rand indices, for example, are used to compare the similarity of two data clusterings. The following section outlines the implementation steps of the NIMAA pipeline and illustrates its application using results from nine diverse datasets analyzed with this methodology.

### 2.1 Bird’s eye view of data structure and submatrix extraction with non-missing values

For a dataset containing two nominal variables, NIMAA constructs a bipartite network. The input data typically consists of a table with two columns representing the nominal variables and an optional third column containing numerical values that serve as edge weights. This information is transformed into an incidence matrix using the plotIncMatrix() function. The incidence matrix is visualized as a heatmap, highlighting both observed and missing values while also capturing structural properties such as modularity and nestedness within the bipartite network. This visualization provides a complementary perspective on the relationships between categories and further illustrates the forms of missingness within the network structure. The resulting matrix object can then be passed to the plotBipartite() function, which generates an interactive igraph-based bipartite network plot, offering a complementary perspective on the relationships and missingness patterns in the data.

In the next step, to extract the complete submatrix with non-missing values from a weighted bipartite network, we use the extractSubMatrix() function. The original matrix is rearranged by this function to find the largest possible submatrix with no missing values. It should be noted that finding the maximal submatrix with no missing values is an NP-hard problem, and this function returns an estimate of the maximal submatrix. For more information on this function, see the NIMAA package vignette.

### 2.2 Analysis of projected networks

As mentioned above, two projected networks are constructed from a bipartite network. The clustering analysis of the projected networks identifies the cluster of similar nodes in each vertex set based on the similarity of neighbors of each node in the bipartite network. The weight of the edges in a weighted bipartite network affects pairwise similarity. With the option to preprocess the input incidence matrix, the findCluster() function generates projected unipartite networks and then applies seven network clustering algorithms. Weak edges can also be removed before clustering. Then, based on internal or external cluster validation indices such as the average silhouette width and Jaccard similarity index, the best clustering method can be chosen. The results are offered as bar plots in addition to a console display. Two external validation indices, the Jaccard similarity coefficient and Rand index, are also provided to measure the similarity between clustering results and the prior knowledge used as ground truth if prior knowledge on the label of nodes is included in the dataset.

### 2.3 Bipartite edge prediction by weight imputation

The predictEdge() function provides an optional downstream analysis step that applies multiple imputation strategies to assess the robustness of structural patterns identified prior to imputation. Rather than treating imputed values as ground truth, this step evaluates whether inferred edges preserve clustering and similarity relationships derived from complete submatrices. Supported techniques include classical approaches such as mean and median imputation, as well as correspondence analysis (CA)^21^ and alternating least squares (ALS)^22^. Once imputation is performed, the resulting completed matrices are projected into unipartite networks for downstream clustering. To evaluate the impact of imputation, the validateEdgePrediction() function compares clustering results obtained from imputed networks against those derived from the maximal submatrix of non-missing values (serving as a benchmark). Several similarity and validation metrics, including the Sørensen-Dice coefficient, Fowlkes-Mallows index, Jaccard similarity, Minkowski index, and Rand index, are computed to quantify the consistency of clustering outcomes^23^. By integrating these metrics, NIMAA enables the systematic selection of the most reliable imputation method, ensuring that edge predictions preserve the underlying structure and relationships within the original nominal dataset.

## 3. Case Studies

In this section, we present nine datasets grouped into four major thematic categories: proteomic data benchmarking, biomedical/pharmacological associations, ecological/environmental networks, and social/behavioral relationships. Each category illustrates specific modules of the nominal data mining workflow, with an emphasis on handling missing values and exploring relationships among nominal labels. The datasets encompass a variety of bipartite network structures, i.e., weighted, unweighted, modular, and nested, demonstrating how the NIMAA pipeline supports pattern recognition, clustering, and predictive modeling across diverse domains.

### 3.1 Proteomic data benchmarking

The bird’s-eye view of a dataset reflects the structure and distribution of missing values and can serve as an important metric for comparing the performance of different analytical tools. This is particularly relevant in proteomic datasets^24^, where a wide variety of software platforms and search engines exist, each with numerous parameters, scoring algorithms, and tuning options. These variations make direct comparisons between tools inherently complex and often inconsistent. To address this challenge, we employ the initial modules of the NIMAA pipeline to assess and compare data completeness and structural integrity in protein quantification outputs by analyzing patterns of missingness. By visualizing and quantifying how missing values are distributed across bipartite representations of the data, NIMAA provides insights into tool-specific biases and structural distortions introduced during processing. We used WOMBAT-P ProteoBenchDDA v0.9.11 protein identification and quantification outputs to conduct a benchmarking pipeline using the NIMAA R package^25^. This framework allows us to evaluate how different quantification tools produce missing data systematically and to identify structural patterns that may affect downstream analyses. These standardized datasets include protein-level outputs from several widely used proteomics software platforms, including MaxQuant, CompOmics, Proline, and TPP.

For consistency across platforms, we extracted columns corresponding to peptide quantification per sample, typically labeled as number_of_peptides_[A/B]_X. Each dataset contained varying numbers of quantified proteins across two conditions, with three replicates each. While the number of columns is fixed across all datasets, the number of quantified proteins (rows) varies. To retain as many proteins as possible while minimizing missing values, we applied the `extractSubMatrix()` function with the mode set to “Rectangular_element_max.” This method identifies the largest possible rectangular submatrix with complete data, maximizing the number of retained proteins (rows) across the fixed set of six samples (columns). Several metrics were then compared, including the size of the non-missing submatrix, nestedness temperature score, relative non-missingness (proportion of observed entries), and overall completeness (proportion of non-missing values in the submatrix relative to the original dataset). These results are summarized in **Tables 1** and **2**, and the row-wise outcome of rectangular max has been visualized in **Fig. 2**.

**Table 1:**
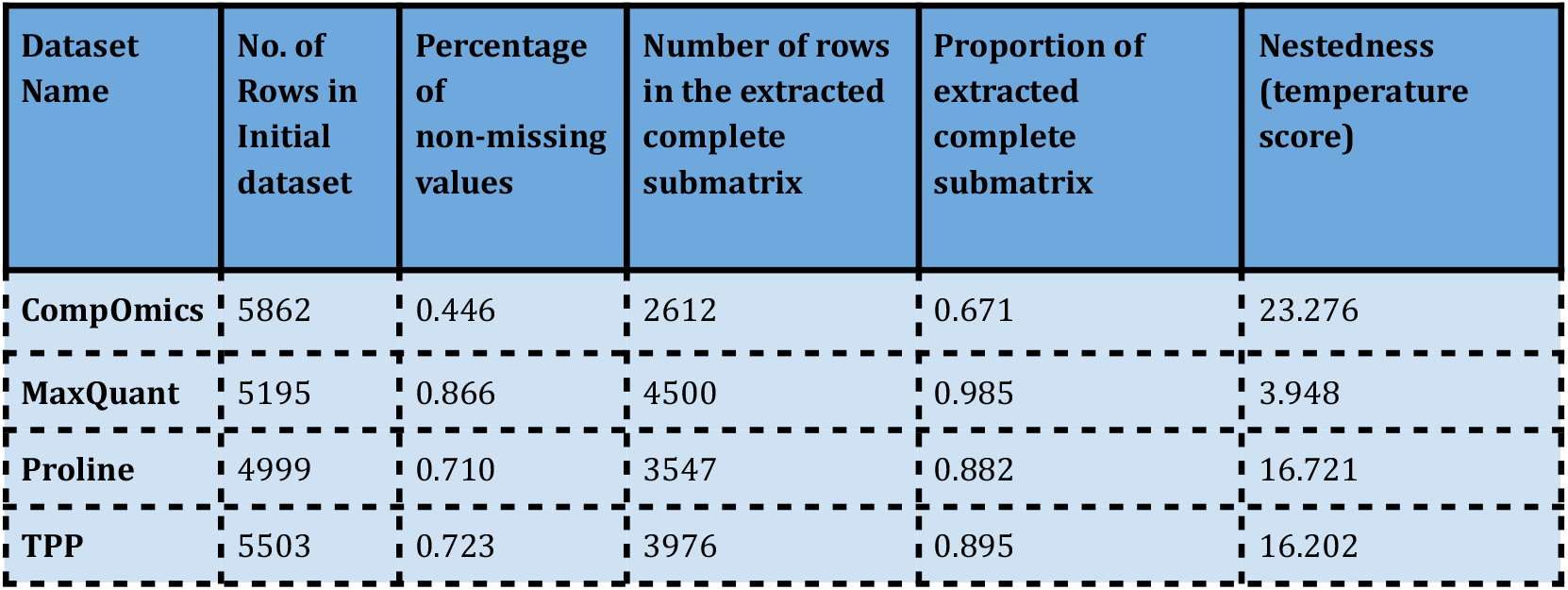
Summary comparison of four proteomics datasets in WOMBAT-P. The table presents the dataset size, the size of the largest complete submatrix, two proportions—(1) total number of non-missing values divided by total possible values (number of rows × 6), and (2) number of rows in the complete submatrix multiplied by 6, divided by total possible values, and the nestedness scores.

**Figure 2.**
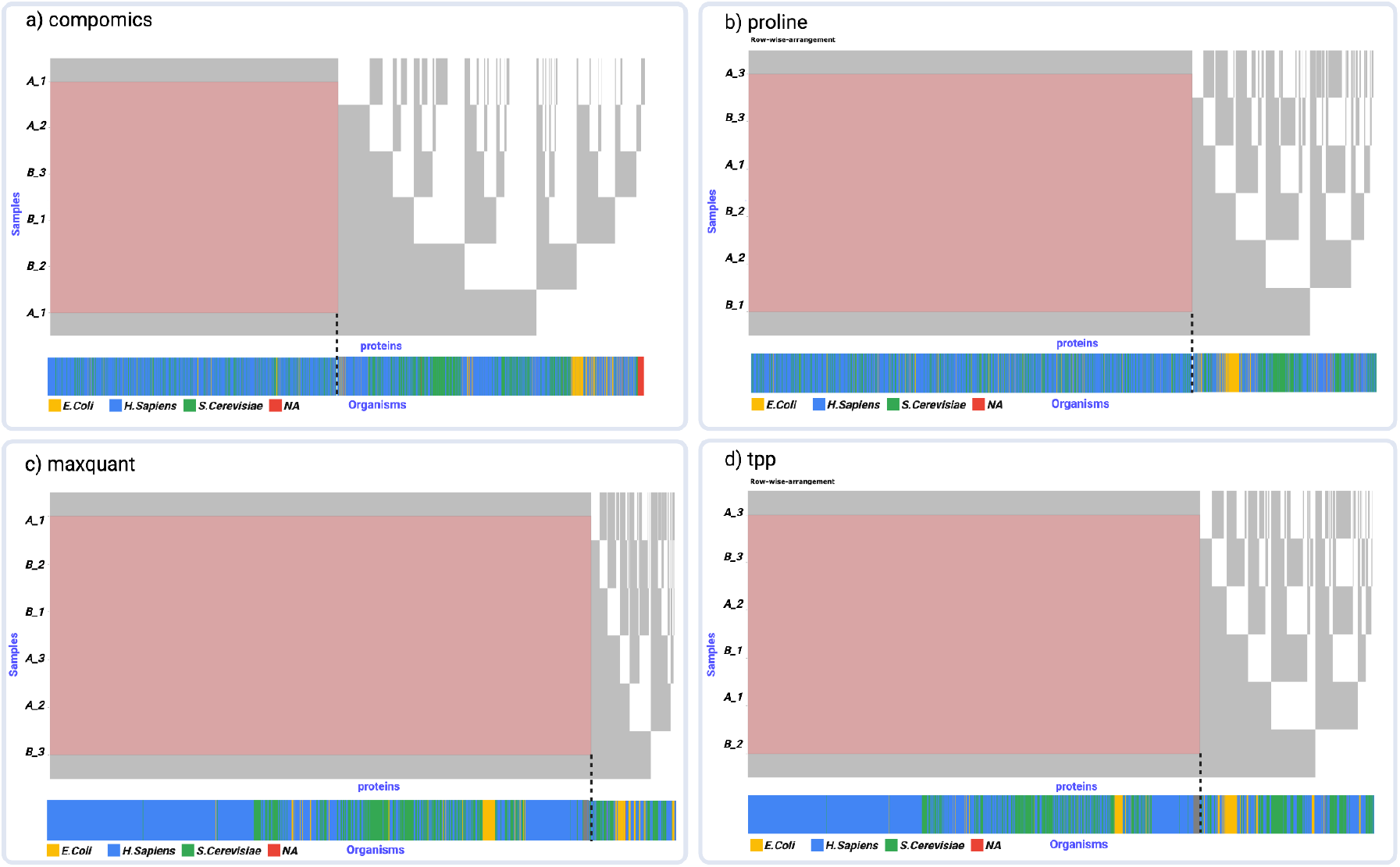
Bird’s-eye view of proteomics datasets. This figure shows the row-wise results of submatrix extraction using the “Rectangular_element_max” mode, applied to each dataset. The method identifies the largest rectangular area without missing values, highlighting the number of complete (non-NA) protein entries retained for each platform: (a) CompOmics, (b) Proline, (c) MaxQuant, and (d) TPP. The heatmap plots next to each dataset represent the species categories, including Homo sapiens (blue), Saccharomyces cerevisiae (green), Escherichia coli (yellow), and NAs (red), which comprise UniParc accession numbers and mostly contain contaminant entries.

MaxQuant demonstrated the highest proportion of extracted complete submatrix (98.5%) and percentage of non-missing values (86.6%), indicating robust and consistent protein quantification. In contrast, CompOmics, although originally reporting the highest number of quantified proteins (5,862), retained the smallest non-missing submatrix and exhibited the lowest nestedness, with a higher temperature score (23.276), suggesting more noise and random missingness. The TPP and Proline platforms showed intermediate performance. Here, performance refers to the ability of each platform to produce protein quantification data that is both complete and structurally coherent, as reflected by the size of the non-missing submatrix, the organization of missing values measured by nestedness and temperature scores, and the biological plausibility of species-level clustering. These platforms had moderate-sized complete submatrices (72.3% and 71%, respectively) and better nestedness scores, reflecting distinct data structures. These findings highlight platform-specific strengths: MaxQuant excels in producing more complete, hierarchically organized data with less noise, while CompOmics shows greater variability and less structured missingness, possibly indicative of MAR (**Fig. 2**). The TPP and Proline outputs fall between these extremes. Such insights are valuable for guiding platform selection based on specific study goals, addressing an ongoing challenge in computational proteomics analysis. Importantly, in this benchmarking setting, all software tools were applied to the same underlying raw mass spectrometry data. Therefore, differences in missingness patterns across platforms do not reflect true biological absence of proteins or differences in detection limits between experiments, but instead arise from tool-specific processing choices, filtering strategies, scoring functions, and identification thresholds. While species-level abundance differences in mixed-species samples provide a biological context, the observed variation in missingness structure primarily captures how different software implementations handle identical input data. In this sense, missing values serve as diagnostic indicators of computational behavior rather than direct biological signals. As shown in **Fig. 2**, this is evident when examining both complete submatrices and those containing missing values. In this figure, the bottom four-color bar (red, blue, green, and grey) denotes proteins originating from *E. coli, H. sapiens, S. cerevisiae*, and proteins of unknown origin, respectively. In the CompOmics and Proline outputs, these colors appear almost randomly distributed, particularly in Nominal data, suggesting minimal organism-specific clustering. In contrast, MaxQuant and TPP exhibit more distinct clustering patterns, with proteins from the same organism grouped together more consistently, indicating a less random and more structured arrangement. Interestingly, we observed that among the benchmarked software packages, MaxQuant outperformed others in terms of the number of identified proteins in the complete submatrix vs. the submatrix with missing values across all organisms. Whilst for TPP and Proline in *E. coli* and for CompOmics in both *E. coli* and yeast, the opposite trend was observed, with more proteins identified in the submatrix containing missing values (**Table 2**). Since human proteins are generally better annotated than those of yeast or *E. coli*, these results indicate that MaxQuant is more effective at handling organisms with less comprehensive annotations, further underscoring its advantage across diverse species. Overall, this comparison underscores that evaluating proteomics software should go beyond simple counts of quantified proteins and instead incorporate completeness, structured missingness, and biological coherence as integrated measures of data quality.

**Table 2:** Species composition in complete and submatrices with missing values of identified proteins across proteomics software outputs. For each dataset generated by CompOmics, MaxQuant, Proline, and TPP, the table reports the frequency of protein identifications per organism within the largest complete submatrix (non-missing values) and in the complementary set containing missing values. Species categories include Homo sapiens, Saccharomyces cerevisiae, Escherichia coli, and NAs, which comprise UniParc accession numbers and mostly contaminant entries.

| Dataset Name | Species-based frequency analysis of complete submatrices/submatrices with missing values |  |  |  |
| --- | --- | --- | --- | --- |
| Species | *Homo sapiens* | *Saccharomyces cerevisiae* | *Escherichia coli* | NAs |
| CompOmics | 1974 / 1576 | 552 / 896 | 57 / 292 | 29 / 18 |
| MaxQuant | 3227 / 303 | 1044 / 246 | 181 / 140 | 48 / 6 |
| Proline | 2608 / 758 | 797 / 472 | 103 / 214 | 39 / 8 |
| TPP | 2913 / 746 | 888 / 532 | 117 / 243 | 58 / 6 |

In addition to completeness and nestedness, we assessed whether the distribution of organism labels within each dataset was random or exhibited clustering patterns using a permutation-based randomness test. This test compared the observed sequence of organisms in each dataset with 5,000 random permutations to derive empirical p-values, indicating whether the observed order was more structured than expected by chance. As summarized in Table 3, both MaxQuant and TPP showed significant non-random clustering (FDR-adjusted p < 0.001) in both complete and missing-value regions, suggesting internally consistent organization and potentially more biologically coherent processing. CompOmics also displayed clustering, though with higher variability between matrix parts, whereas Proline’s complete submatrix was indistinguishable from random (p_adj ≈ 0.21), implying that its filtering step reduces organism-specific bias. Together, these results reinforce that the missingness structure itself encodes information about software-specific data organization.

**Table 3:** Empirical permutation tests of organism-ordering randomness. Lower adjusted p-values indicate that the observed order of organism labels is less random than expected under permutation, reflecting structured clustering. Consistent clustering in both complete and missing regions (as in MaxQuant and TPP) suggests systematic organization of quantification results, while a random pattern (as in Proline’s complete submatrix) indicates unbiased distribution after filtering.

| Software | Matrix part | p_adj | Sig | Interpretation |
| --- | --- | --- | --- | --- |
| CompOmics | dropped (missing part) | 0.00027 | *** | non-random (clustered) |
| CompOmics | kept (complete submatrix) | 0.0103 | * | non-random (clustered) |
| MaxQuant | dropped (missing part) | 0.00027 | *** | non-random (clustered) |
| MaxQuant | kept (complete submatrix) | 0.00027 | *** | non-random<br>(clustered) |
| Proline | dropped (missing part) | 0.00027 | *** | non-random<br>(clustered) |
| Proline | kept (complete submatrix) | 0.215 | ns | indistinguishable<br>from random |
| TPP | dropped (missing part) | 0.00027 | *** | non-random<br>(clustered) |
| TPP | kept (complete submatrix) | 0.00027 | *** | non-random<br>(clustered) |

### 3.2 Biomedical and Pharmacological Networks

We applied the NIMAA framework to three diverse datasets; BeatAML, Herb-Ingredient, and DrugComb, each representing distinct biomedical domains. These datasets vary in scale, structure, and missingness patterns, offering complementary perspectives for evaluating the utility of NIMAA workflow analysis.

#### 3.2.1 beatAML Dataset

The BeatAML dataset contains drug sensitivity measurements from *ex vivo* acute myeloid leukemia (AML) patient samples, offering a valuable resource for evaluating the efficacy of different inhibitors ^26^. It comprises 122 distinct inhibitors tested across 528 patient samples and can be represented as a bipartite network, where the two nominal variables are inhibitors and patient identifiers. Edge weights correspond to the median drug response values, enabling detailed comparisons of drug sensitivity patterns across patients. Analysis of the incidence matrix reveals a nestedness score of 20.12, reflecting a moderate hierarchical structure in drug–patient associations. The dataset contains 26% missing values, with the largest square complete submatrix (free of missing entries) covering 96 inhibitors and 96 patients (**Fig. 3, panel a**). These missing values are not randomly distributed; instead, their structure encodes meaningful constraints in the experimental space.

**Figure 3.**
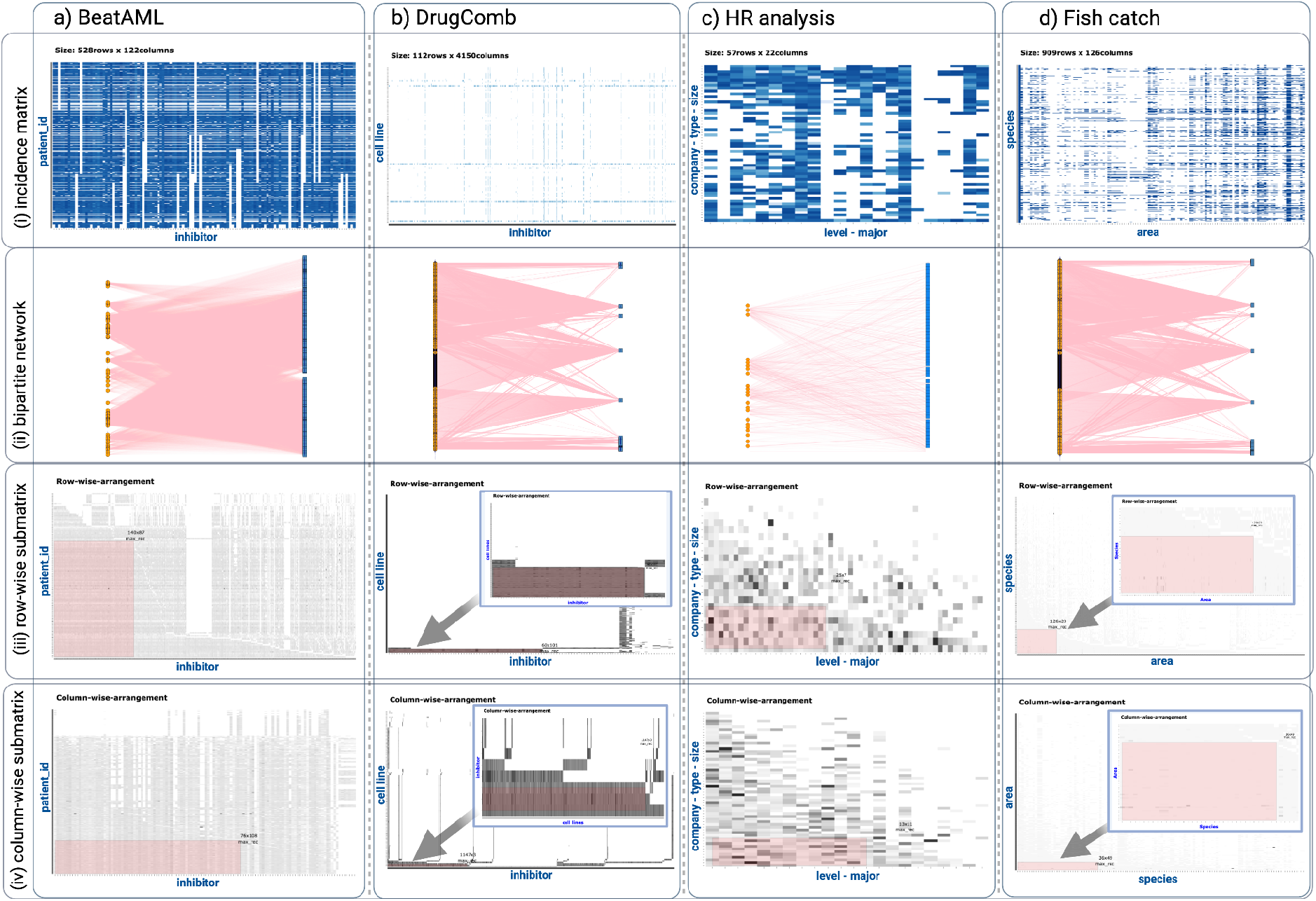
Visualizing data completeness and structure across diverse datasets using NIMAA. For each dataset (a) BeatAML drug response, (b) DrugComb synergy analysis, (c) HR analytics, and (d) Fish catch records, the panels display (i) the incidence matrix highlighting the overall proportion of missing values, (ii) the corresponding bipartite network of nominal variables, (iii) the extracted row-wise submatrix with non-missing values, and (iv) the extracted column-wise submatrix with non-missing values.

Previous work by Jafari et al. ^27^ demonstrated how leveraging the topology of missingness in such bipartite networks, specifically by examining the complement space of absent associations, can uncover novel and clinically relevant drug combination strategies for AML. In this context, the missing value structure itself becomes an informative layer, guiding the identification of promising inhibitor–patient pairings for combination therapy design.

#### 3.2.2 DrugComb dataset

The DrugComb dataset serves as a comprehensive resource that integrates cancer drug combination screening studies from various high-throughput experiments ^28^. It provides data on synergistic and antagonistic effects of drug pairs tested across multiple cancer cell lines. In the NIMAA framework, this dataset is structured as a bipartite network in which the nominal variables represent drug combinations and corresponding cell lines. The edge weights reflect the observed synergy scores, such as the Bliss or Loewe models. This configuration facilitates an in-depth exploration of combinatorial effects and potential relationships between drug pairs and specific cancer types (see the NIMAA Shiny app for details). Missing synergy values are imputed and analyzed using clustering and projection methods to uncover latent patterns and promising combinations for follow-up studies. The resulting patterns help prioritize promising combinations, enabling the detection of therapeutic synergies and cell line-specific responses.

#### 3.2.3 Herb-Ingredient dataset

The second version of the TCMID (Traditional Chinese Medicine Integrated Database) is a key resource in the field of Traditional Chinese Medicine (TCM) herbs ^29^. TCMID provided a comprehensive resource by incorporating ingredient-specific experimental data obtained from herbal mass spectrometry spectra. This extensive dataset encompassed 2,815 ingredients and 3,775 herbs. To ensure an adequate amount of data, a filtering criterion was applied, retaining only those herbs that contained a minimum of 40 ingredients and ingredients that were present in at least 40 herbs. Consequently, a dataset comprising 51 ingredients and 28 herbs was obtained (see the NIMAA Shiny app for details). This heterogeneous nature of herbal formulations suggests that ingredient sharing across herbs may be driven more by empirical tradition than systematic design.

Building on such ingredient–herb associations, earlier work by Jafari et al. ^15^ demonstrated how bipartite network analysis of ingredient–herb relationships in TCM can reveal molecular patterns underlying traditional classifications, such as meridians and properties. In this context, the alignment between ingredient clusters and classification profiles becomes an informative layer, bridging empirical TCM knowledge with modern chemical and molecular data to support phenotype-based drug discovery.

### 3.3 Ecological and Environmental Networks

To demonstrate NIMAA’s applicability beyond biomedical contexts, we analyzed two datasets from ecology and environmental monitoring: the historical Robertson plant–pollinator network and the Fish Catch Annual dataset. These datasets illustrate how structured missingness can capture ecological specialization, temporal dynamics, and environmental constraints, providing insights that go beyond observed interactions or measurements.

#### 3.3.1 Robertson data set

Charles Robertson’s historical plant–pollinator dataset provides a unique opportunity to study the structure of missing interactions over an extended timescale. Collected between 1884 and 1916 in Macoupin County, Illinois, USA, the data originally recorded 1,429 insect species visiting 456 plant species. When represented as a bipartite incidence matrix, missing values correspond to unobserved or absent interactions between specific plant–pollinator pairs ^30^.

To improve interpretability, we applied a filtering step, retaining only insects that visited at least 30 flowers and plants visited by at least 50 insects. This yielded a reduced network of 122 pollinator species and 24 plant species. The resulting matrix reveals distinct patterns of missingness: clusters of complete interaction data for some pollinator–plant groups coexist with large, structured gaps for others (see the NIMAA Shiny app for details). For example, eight pollinator species form complete interaction blocks with eight plant species (specialized relationships), while 24 pollinators interact with only four plants, generating a more sparsely populated yet patterned submatrix (generalized relationships). These non-random missingness structures reflect ecological realities, such as specialization, seasonal overlap, and spatial constraints, rather than mere absence of sampling, demonstrating how structured gaps themselves can encode functional roles and community organization in a stable but complex ecosystem.

#### 3.3.2 Fish Catch Annual Data Set

The fish catch annual dataset documents annual fish and shellfish catches in the Northeast Atlantic across multiple decades, where rows represent species and columns represent areas. Missing values in the bipartite incidence matrix correspond to zero-reported or unrecorded catches for certain area–species combinations over 6 years (2009-2014). Visual inspection and nestedness analysis, with the score of 2.66, reveal that these gaps are not uniformly distributed: some species exhibit continuous reporting across years in specific areas, while others display patchy but temporally clustered absences. Such structured missingness may reflect historical changes in fishing effort ^31^, climate change-related species availability ^32^, or regulatory mechanisms ^33^, rather than random omissions.

By projecting the network and clustering species or years, NIMAA highlights co-varying groups where missing values align across blocks, suggesting shared ecological or management drivers (**Fig. 3, Panel d**). This structured gap patterning provides additional insight into long-term biodiversity changes and fishing practices, reinforcing the importance of interpreting the missingness form alongside observed values in sustainability analyses.

### 3.4 Human Behavior and Social Systems

We further applied NIMAA to datasets from human resources, online education, and retail transactions, demonstrating its versatility across social and commercial domains. In each case, structured missingness revealed meaningful subgroup patterns and behavioral signals, offering insights that extend beyond the recorded data.

#### 3.4.1 Human Resource (HR) Analytics dataset

In the HR analytics dataset, employees and organizational attributes (e.g., department, job satisfaction, promotion status) form the two nominal variable sets. Missing entries, such as absent performance ratings, do not occur randomly but often cluster within specific employee groups or organizational units. This block-like missingness may arise from policy differences, survey non-response patterns, or role-specific reporting gaps.

When visualized as an incidence matrix, these missing clusters can signal latent subgroups with shared work environments or cultural dynamics (**Fig. 3, Panel c**). Instead of immediate imputation, NIMAA first leverages these patterns to guide clustering, revealing associations between structured missingness and turnover risk. This approach ensures that missing values contribute to the understanding of organizational behavior rather than being treated solely as noise.

#### 3.4.2 Student Discussion data set

The student discussion dataset models students and discussion topics from an e-learning platform as a bipartite graph. Missing values correspond to unengaged topic–student pairs. Far from being random, these gaps often form topic-specific blocks, where certain groups of students consistently avoid or remain inactive in particular threads. Such structured absence patterns can indicate topical disengagement, prerequisite knowledge gaps, or social clustering in participation.

By analyzing the topology of these gaps before imputation, NIMAA helps identify passive learner communities and isolated topics, allowing targeted interventions (see the NIMAA Shiny app for details). Structured missingness here becomes an educational signal, enabling the detection of learning bottlenecks and informing personalized course design.

#### 3.4.3 Online Retail data set

Here in the retail transaction dataset, customers and products form a bipartite network where edge weights reflect purchase volume or value. Missing values dominate the incidence matrix, representing customer–product pairs with no recorded purchases. These absences are not uniformly distributed: certain product groups show high co-purchase coverage among specific customer clusters, while others are completely absent for those same groups, forming structured “void” regions in the matrix.

Such patterns often reflect product specialization, seasonality, or targeted marketing effects. NIMAA’s analysis reveals how these structured absences can complement purchase data to refine customer segmentation and product recommendation strategies (see the NIMAA Shiny app for details). By preserving the shape of missingness in early analysis, retailers can detect latent demand opportunities and optimize recommendation systems with more nuanced market insights.

## 4. Conclusion

This work reframes missing value forms in nominal datasets from being mere analytical hindrances to becoming informative structural features. By modeling data as bipartite networks and examining the form, distribution, and topology of missingness, the NIMAA framework reveals that gaps often encode latent biological constraints, operational biases, or domain-specific organization. Across proteomics benchmarking, pharmacogenomics, ecological surveys, and social systems, we demonstrate that structured missingness can illuminate system modularity, nestedness, and category-level similarities, often in ways that enhance interpretability beyond what observed values alone can provide. Rather than defaulting to early imputation or exclusion, our approach prioritizes the analysis of missingness structure before any value inference, using optional imputation only as a secondary step to assess the stability of the identified patterns.. These findings advocate for a paradigm shift: missing values should be treated not simply as noise to be corrected, but as valuable signals capable of enriching discovery and guiding more context-aware modeling in complex nominal systems.

## Data and code availability

All data sources used in this study are linked to their corresponding panels in the Shiny web application, available at https://ehsan-zangene.shinyapps.io/nimaa_app/. The analysis codes are publicly accessible at https://github.com/esnzgn/NIMAA_cases. In particular, the benchmarking code and high-resolution figure outputs for the NIMAA mass spectrometry proteomics analysis are available at https://github.com/esnzgn/NIMAA_MS_bench. The NIMAA R package itself is available through the CRAN repository.

## References

1. Pham, T. M., Pandis, N. & White, I. R. Missing data, part 1. Why missing data are a problem. Am J Orthod Dentofacial Orthop 161, 888–889 (2022).

2. Pham, T. M., Pandis, N. & White, I. R. Missing data, part 2. Missing data mechanisms: Missing completely at random, missing at random, missing not at random, and why they matter. Am J Orthod Dentofacial Orthop 162, 138–139 (2022).

3. Panda, B. S. & Kumar Adhikari, R. A method for classification of missing values using data mining techniques. In 2020 International Conference on Computer Science, Engineering and Applications (ICCSEA) (IEEE, 2020). doi:10.1109/iccsea49143.2020.9132935.

4. Little, R. J. Missing data assumptions. Annu. Rev. Stat. Appl. 8, 89–107 (2021).

5. Little, R. J. Missing Data Analysis. Annu Rev Clin Psychol 20, 149–173 (2024).

6. García-Laencina, P. J., Sancho-Gómez, J.-L. & Figueiras-Vidal, A. R. Pattern classification with missing data: a review. Neural Comput. Appl. 19, 263–282 (2010).

7. Bhinder, B., Gilvary, C., Madhukar, N. S. & Elemento, O. Artificial Intelligence in Cancer Research and Precision Medicine. Cancer Discov 11, 900–915 (2021).

8. Josephson, C. B. & Wiebe, S. Precision Medicine: Academic dreaming or clinical reality? Epilepsia 62 Suppl 2, S78–S89 (2021).

9. Jafari, M., Guan, Y., Wedge, D. C. & Ansari-Pour, N. Re-evaluating experimental validation in the Big Data Era: a conceptual argument. Genome Biol 22, 71 (2021).

10. Zangene, E.Marashi, S.-A. & Montazeri, H. SL-scan identifies synthetic lethal interactions in cancer using metabolic networks. Sci Rep 13, 15763 (2023).

11. Siedlecki, S. L. & Bena, J. F. Exploring Associations When Your Data are Nominal: A Review of the χ2 Test. Clin Nurse Spec 35, 229–232 (2021).

12. Suich, R. & Turek, R. Intuition in using nominal variables for prediction. Teach. Stat. 25, 86–89 (2003).

13. Che, Z., Purushotham, S., Cho, K., Sontag, D. & Liu, Y. Recurrent Neural Networks for Multivariate Time Series with Missing Values. Sci Rep 8, 6085 (2018).

14. Turanli, B. et al. A Network-Based Cancer Drug Discovery: From Integrated Multi-Omics Approaches to Precision Medicine. Curr Pharm Des 24, 3778–3790 (2018).

15. Jafari, M., Wang, Y., Amiryousefi, A. & Tang, J. Unsupervised Learning and Multipartite Network Models: A Promising Approach for Understanding Traditional Medicine. Front Pharmacol 11, 1319 (2020).

16. Nominal Data Mining Analysis [R package NIMAA version 0.2.1]. (2022).

17. Pavlopoulos, G. A. et al. Bipartite graphs in systems biology and medicine: a survey of methods and applications. Gigascience 7, 1–31 (2018).

18. Patterson, B. D. & Atmar, W. Nested subsets and the structure of insular mammalian faunas and archipelagos. Biol. J. Linn. Soc. Lond. 28, 65–82 (1986).

19. Jafari, M., Sadeghi, M., Mirzaie, M.Marashi, S.-A. & Rezaei-Tavirani, M. Evolutionarily conserved motifs and modules in mitochondrial protein-protein interaction networks. Mitochondrion 13, 668–675 (2013).

20. Bartholdi, J. J., III. A good submatrix is hard to find. Oper. Res. Lett. 1, 190–193 (1982).

21. Josse, J. & Husson, F. MissMDA: A package for handling missing values in multivariate data analysis. J. Stat. Softw. 70, (2016).

22. Hastie, T. & Mazumder, R. softImpute: Matrix Completion via Iterative Soft-Thresholded SVD. CRAN: Contributed Packages The R Foundation 10.32614/cran.package.softimpute (2013).

23. Jafari, M., Mirzaie, M. & Sadeghi, M. Interlog protein network: an evolutionary benchmark of protein interaction networks for the evaluation of clustering algorithms. BMC Bioinformatics 16, 319 (2015).

24. Lazar, C., Gatto, L., Ferro, M., Bruley, C. & Burger, T. Accounting for the Multiple Natures of Missing Values in Label-Free Quantitative Proteomics Data Sets to Compare Imputation Strategies. J Proteome Res 15, 1116–1125 (2016).

25. Bouyssié, D. et al. WOMBAT-P: Benchmarking Label-Free Proteomics Data Analysis Workflows. Journal of Proteome Research (2023) doi:10.1101/2023.10.02.560412.

26. Tyner, J. W. et al. Functional genomic landscape of acute myeloid leukaemia. Nature 562, 526–531 (2018).

27. Jafari, M. et al. Bipartite network models to design combination therapies in acute myeloid leukaemia. Nat Commun 13, 2128 (2022).

28. Zagidullin, B. et al. DrugComb: an integrative cancer drug combination data portal. Nucleic Acids Res 47, W43–W51 (2019).

29. Huang, L. et al. TCMID 2.0: a comprehensive resource for TCM. Nucleic Acids Res 46, D1117–D1120 (2018).

30. Marlin, J. C. & Laberge, W. The native bee fauna of Carlinville, Illinois, revisited after 75 years: A case for persistence. Conservation Ecology 5, 9 (2001).

31. Barcellos, L. R., da Silva, T. E. F. & Lessa, R. P. T. Historical changes in conservation measurements on target, predictable bycatch and bycatch species caught by pelagic longline fisheries in Southwest Atlantic Ocean. Mar. Policy 173, 106567 (2025).

32. Bastardie, F. et al. Anticipating how spatial fishing restrictions in EU waters perform to protect marine species, habitats, and dependent fisheries. Front. Mar. Sci. 12, (2025).

33. Harte, M., Tiller, R., Kailis, G. & Burden, M. Countering a climate of instability: the future of relative stability under the Common Fisheries Policy. ICES J. Mar. Sci. 76, 1951–1958 (2019).

